# MemBrain: A Deep Learning-aided Pipeline for Automated Detection of Membrane Proteins in Cryo-electron Tomograms

**DOI:** 10.1101/2022.03.01.480844

**Authors:** Lorenz Lamm, Ricardo D. Righetto, Wojciech Wietrzynski, Matthias Pöge, Antonio Martinez-Sanchez, Tingying Peng, Benjamin D. Engel

**Affiliations:** Helmholtz Pioneer Campus, Helmholtz Munich, 85764 Neuherberg, Germany; Helmholtz AI, Helmholtz Munich, 85764 Neuherberg, Germany; iozentrum, University of Basel, 4056 Basel, Switzerland; Max Planck Institute of Biochemistry, 82152 Martinsried, Germany; Department of Computer Science, Faculty of Sciences - Campus Llamaquique, University of Oviedo, 33007 Oviedo, Spain; Health Research Institute of Asturias (ISPA), Avenida Hospital Universitario s/n, 33011 Oviedo, Spain

**Keywords:** Cryo-electron tomography, particle picking, protein localization, membrane protein, deep learning, label-efficient

## Abstract

**Background and Objective:** Cryo-electron tomography (cryo-ET) is an imaging technique that enables 3D visualization of the native cellular environment at sub-nanometer resolution, providing unpreceded insights into the molecular organization of cells. However, cryo-electron tomograms suffer from low signal-to-noise ratios and anisotropic resolution, which makes subsequent image analysis challenging. In particular, the automated detection of membrane-embedded proteins is a problem still lacking satisfactory solutions.

**Methods:** We present MemBrain – a new deep learning-based pipeline that automatically detects membrane-bound protein complexes in cryo-electron tomograms. After subvolumes are sampled along a segmented membrane, each subvolume is assigned a score using a convolutional neural network (CNN), and protein positions are extracted by a clustering algorithm. Incorporating rotational subvolume normalization and using a tiny receptive field simplify the task of protein detection and thus facilitate the network training.

**Results:** MemBrain requires only a small quantity of training labels and achieves excellent performance with only a single annotated membrane (F1 score: 0.88). A detailed evaluation shows that our fully trained pipeline outperforms existing classical computer vision-based and CNN-based approaches by a large margin (F1 score: 0.92 vs. max. 0.63). Furthermore, in addition to protein center positions, MemBrain can determine protein orientations, which has not been implemented by any existing CNN-based method to date. We also show that a pre-trained MemBrain program generalizes to tomograms acquired using different cryo-ET methods and depicting different types of cells.

**Conclusions:** MemBrain is a powerful and label-efficient tool for the detection of membrane protein complexes in cryo-ET data, with the potential to be used in a wide range of biological studies. It is generalizable to various kinds of tomograms, making it possible to use pretrained models for different tasks. Its efficiency in terms of required annotations also allows rapid training and fine-tuning of models. The corresponding code, pretrained models, and instructions for operating the MemBrain program can be found at: https://github.com/CellArchLab/MemBrain

## 1 Introduction

Cryo-electron tomography (cryo-ET) is performed by acquiring projection images of a frozen sample at numerous different angles using an electron microscope. Combining the information from all these images yields high-resolution 3D volumes. These molecular views provide a striking new perspective on cellular functions and can reveal important insights into both general biological phenomena [1] as well as human diseases including neurodegeneration and cancer [2,3]. About one third of human proteins are associated with membranes [4], which compartmentalize the cell. Membranes organize proteins into microdomains to direct cellular processes, including the bioenergetic reactions in mitochondria and chloroplasts that produce the energy of life. It is thus an important challenge to identify protein complexes within their native membrane environment.

In cryo-ET, particle picking is commonly performed using template matching [5], i.e., a low-pass filtered model of the macromolecular complex of interest is fit to the entire tomogram by computing cross-correlation scores. Since this has to be done for all possible positions and orientations, this approach is computationally expensive and yields a noisy output that requires manual post-processing. For membrane-bound proteins, template matching is particularly challenging, as these complexes are often small, and large portions of their structures are buried within the membranes, which can reduce their correlation scores with the template.

Besides template matching, PySeg [6] is a template-free procedure for the detection of membrane-bound proteins. It uses discrete Morse theory to trace densities protruding from the membrane surface. These densities are then clustered using 2D rotational averages, followed by manual selection of reasonable classes. However, the use of PySeg is complicated by difficult parameter tuning and the requirement for human intervention.

Recently, several deep learning-based approaches have also been proposed for the detection of macromolecular complexes. EMAN [7] detects various kinds of densities by creating segmentations based on a 2D CNN architecture that utilizes large convolutional kernels of size 15×15. Therefore, it requires large image patches for training, where all present objects of interest must be annotated. This is also the case for DeepFinder [8], which requires even more annotations, since it utilizes 3D convolutions with a large receptive field. Another approach is implemented in crYOLO [9], which was initially developed for detecting single particles in 2D cryo-EM micrographs, but has since added the option to pick in 3D tomograms. It is based on the popular object detection framework YOLO [10]. Instead of creating a segmentation mask first, crYOLO predicts the bounding boxes of the particles directly. An advantage of crYOLO is that it requires only positive labels and uses adjustable penalty weights for false negatives and false positives to account for incompletely annotated regions. Nevertheless, we experienced that crYOLO fails to perform reliable predictions when trained with sparse annotations.

In current cryo-ET analysis workflows, manual particle picking is often performed to localize membrane-bound proteins. Even with the guidance of specialized software [11,12], the accurate annotation of particles in 3D remains a tedious task. Moreover, tomograms suffer from a low signal-to-noise ratio and anisotropic resolution from the missing wedge [13]. Thus, precisely localizing protein complexes remains a major challenge with existing methods, especially in large tomograms that involve thousands of proteins of interest. To enable new biological discoveries, it is necessary to develop automated tools that can be trained on a small number of high-quality annotations to reliably pick particles in extensive datasets of many tomograms.

Here, we present MemBrain: a label-efficient, generalizable deep learning-based pipeline to detect membrane-bound protein complexes in cryo-ET. In contrast to other approaches, MemBrain utilizes the membrane geometry to sample and align subvolumes and thereby achieve model invariance to unseen membrane orientations. Furthermore, instead of classifying each subvolume into binary classes – i.e., particle vs. non-particle – we show that score assignment based on the regression of distance to the nearest particle can cope better with imprecise ground-truth positions, which commonly occur in manual labelling. Additionally, MemBrain can extract protein orientations, as compared to common detection networks that only provide protein positions. To our knowledge, this is the first deep learning-based pipeline that is specialized for the detection of membrane-bound proteins.

## 2 Methodology

### 2.1 Overview of MemBrain

Figure 1 illustrates the workflow of our pipeline. As an input, we use membrane segmentations from raw cryo-electron tomograms (Fig. 1A). For example, these segmentations can be generated using TomoSegMemTV [14], followed by manual curation, as demonstrated in [12]. In the first step, we sample points uniformly on the segmented membrane and identify their corresponding membrane normal vectors. Subsequently, subvolumes are extracted around each sampled position and rotated so that the membrane is parallel to the x-y plane (pre-processing step, Fig. 1B). These aligned subvolumes are then fed into a CNN, which assigns each subvolume a score indicating its distance to a particle center (scoring step, Fig. 1C). Finally, particle center positions and orientations are extracted using Mean Shift clustering [15] (post-processing step, Fig. 1D). We provide a detailed explanation of each step below.

**Fig. 1.**
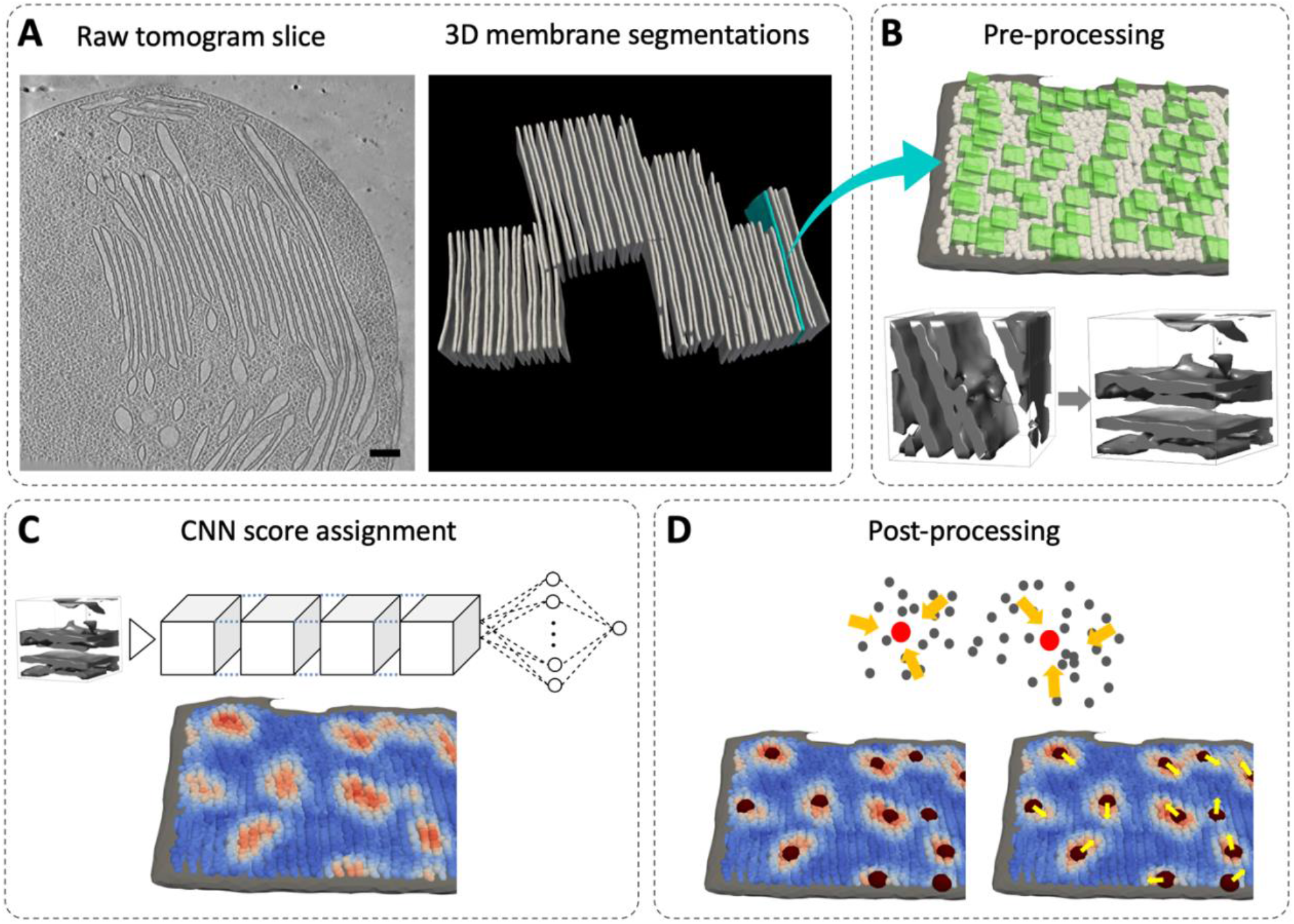
The MemBrain processing pipeline. **A:** An example 2D slice of a cryo-electron tomogram (denoised using Cryo-CARE [25] scale bar 100nm) and corresponding membrane segmentations (3D view of all segmented membranes). **B:** Pre-processing step: Top: Points (white) are sampled and subvolumes (green) around each of them are extracted. Bottom: Subvolumes are aligned. **C:** Scoring step: CNN predicts the distance of each subvolume to the nearest particle center. **D:** Post-processing: particle centers and orientations are extracted using Mean Shift Clustering.

### 2.2 Subvolume Sampling and Preprocessing

#### Point Sampling and Normal Voting

We first manually specify which side of the membrane should be picked. This is done by clicking on the desired side of the membrane segmentation (illustrated in Fig. S6 A). Next, MemBrain samples points all over the selected membrane surface (see sampled positions in Figure 1B). More specifically, we compute the distance of any voxel in the tomogram to the membrane and threshold this distance to mask the neighborhood of the segmentation (see Fig. S6 B; in all our experiments, we used a maximum distance of 15 voxels). Then, for each point in this neighborhood, we find its nearest voxel on the segmented membrane surface. These points on the surface represent our sampled points. The connection vector between them and the original points in the neighborhood serve as a membrane normal vector.

However, points in the membrane neighborhood that are very close to the surface tend to have non-reliable normal vectors. On the other hand, only considering points with large distances leads to non-uniform point sampling on the membrane surface. Therefore, we apply Normal Voting [16,17], which is an algorithm that corrects normal vectors via weighted averaging, while also taking the membrane curvature into account. By giving a higher weight to the reliable normal vectors (from long-distance points), we achieve both a uniform sampling of points on the membrane surface and a reliable estimation of normal vectors.

#### Subvolume Extraction and Rotation

For each sampled position on the membrane surface, we extract a small subvolume. The exact size of the subvolume should depend on the size of the protein complex we aim to detect. In our experiments, we use a subvolume size of 12×12×12. We experienced this size to be quite robust to different protein dimensions, but it can be adjusted if needed. Next, we rotate each subvolume using its corresponding normal vector so that the membrane contained in the subvolume is aligned to the x-y plane, i.e., the membrane normal vector is parallel to the z-axis (see rotated volume in Figure 1B). This rotation step is critical to guarantee that our detection is invariant to different membrane orientations, as we elaborate in Section 3.

### 2.3 Convolutional Neural Network

#### Architecture

The preprocessing steps of MemBrain reduce the task of protein detection in the entire tomogram to the detection of densities protruding from the membrane. Since the context, i.e., the location of the membrane, is already given by the membrane segmentation, our network does not need to learn it. Therefore, we can neglect large-scale features and use a small network architecture, resulting in a tiny receptive field of the convolutional layers. In our experiments, we only used four consecutive 3D convolutional layers, followed by a single fully connected layer. The network outputs a single value to indicate the distance of the subvolume to the nearest protein center. For the detailed network architecture, see Table S4.

#### Label Assignment

For training, we assigned labels to each input subvolume based on its center point’s distance to the closest protein complex center. However, protein structures are generally not spherical. Therefore, instead of using the Euclidean distance to assign the distance values, we use the Mahalanobis distance [18], which weights distances in each direction based on a covariance matrix, thus taking protein shape into account (visualized in Figure S8). This covariance matrix can be extracted from a low-pass filtered structure of respective protein.

#### Optimization Details

Even with substantial effort in manual annotation, our ground-truth labels are not perfectly accurate, as manually picked positions always deviate slightly from the exact particle locations. Moreover, it is almost unavoidable to miss some protein complexes. Therefore, we choose to optimize the smooth L1 loss function [19], which is more robust with respect to outliers compared to the commonly used mean squared error (MSE). We use the Adam [20] optimizer to minimize the loss function. To improve training stability, we incorporate several batch normalization layers [21]. Finally, as regularization techniques, we use a combination of weight decay and data augmentation. In particular, we use random rotations around the z-axis, random flipping along the x- and y-axes to artificially increase the number of samples, as well as adding Gaussian noise to the subvolumes [22].

### 2.4 Post-processing

#### Mean Shift Clustering

We utilize the distance heatmap that results from our network predictions to extract particle positions. For this, we use only the points with a predicted distance less than a threshold (depending on the protein complex size) to perform our adapted version of Mean Shift clustering, where we have added a hierarchical cluster separation step. Mean Shift [15] is an iterative approach that refines cluster centers based on a moving weighted average in each iteration. The only required parameter is the bandwidth, which specifies the radius considered for the weighted average. Using a large bandwidth leads to particle clusters that are merged, whereas a small bandwidth can cause one particle to split into two clusters. To avoid both of these errors, we implemented a refined version of Mean Shift: first, we choose a large bandwidth parameter, leading to relatively large clusters. If one of these clusters exceeds a certain size (depending on the protein complex size), we reprocess it using a lower bandwidth. Thereby, we split merged protein clusters hierarchically. Our version of Mean Shift also weights single points using the inverse network-assigned scores to guide the clustering algorithm towards the correct particle centers, since points close to them tend to have stronger (i.e., lower) distance values.

#### Orientation Estimation

Besides the cluster center, Mean Shift also identifies each point that has converged to the respective cluster center. We assume that these points resemble the rough shapes of the protein complexes. Thus, in case of non-spherical proteins, we can use the cluster points to compute orientations of the detected protein complexes within the membrane. We perform Principal Component Analysis [23] to find the vector that best describes the cluster, i.e., minimizes the Euclidean distances between the cluster points and the line described by the vector. In combination with the previously extracted normal vector, we can compute all three Euler angles describing the protein’s orientation in space. These Euler angles can be used to initialize downstream tasks, such as subtomogram averaging [13].

## 3 Results

### 3.1. Data Collection and Annotation

We evaluated MemBrain using three cryo-ET datasets of different biological samples that were vitrified on EM grids and then thinned with focused ion beam (FIB) milling [24]. Dataset 1 consists of nine tomograms of isolated *Spinacia oleracea* chloroplasts, acquired with defocus imaging and denoised with the deep learning-based Cryo-CARE program [25]. Dataset 2 [12] contains four tomograms of *Chlamydomonas reinhardtii* chloroplasts inside native cells, acquired with Volta phase plate (VPP) imaging and not denoised. Dataset 3 [26] is composed of five tomograms of isolated rod outer segments from wild-type mice, acquired without VPP and denoised with Cryo-CARE. Figure 2 shows example 2D slices of all three datasets, as well as Membranogram views that display the different distributions of protein complexes within the membranes. More detailed information about acquisition of the dataset can be found in the supplementary S.1.1. In all tomograms, membranes were segmented using TomoSegMemTV [14], followed by manual curation. For Datasets 1 and 2, particle positions and orientations were manually annotated using Membranorama [11,12], whereas for Dataset 3, particle positions were generated semiautomatically by an expert using PySeg (see supplementary S.1.2). For Dataset 1, a total of 455 membrane segmentations have been created, 45 of which are annotated with protein complex locations and orientations for 1641 Photosystem II (PSII), 471 Cytochrome b6f (b6f) proteins, and 757 unknown densities (UK). Dataset 2 contains a total of 31 segmented and annotated membranes, including 730 PSII positions, 379 b6f positions and 273 UK positions. For training and validation sets, we used 35 membranes from seven Dataset 1 tomograms. We trained and evaluated with PSII complexes, as the b6f complexes are so small that they are hard to visually identify in some tomograms, and therefore will require further adjustments to reliably detect. The remaining two tomograms from Dataset 1, as well as all tomograms from Datasets 2 and 3, were reserved as test sets.

**Fig. 2.**
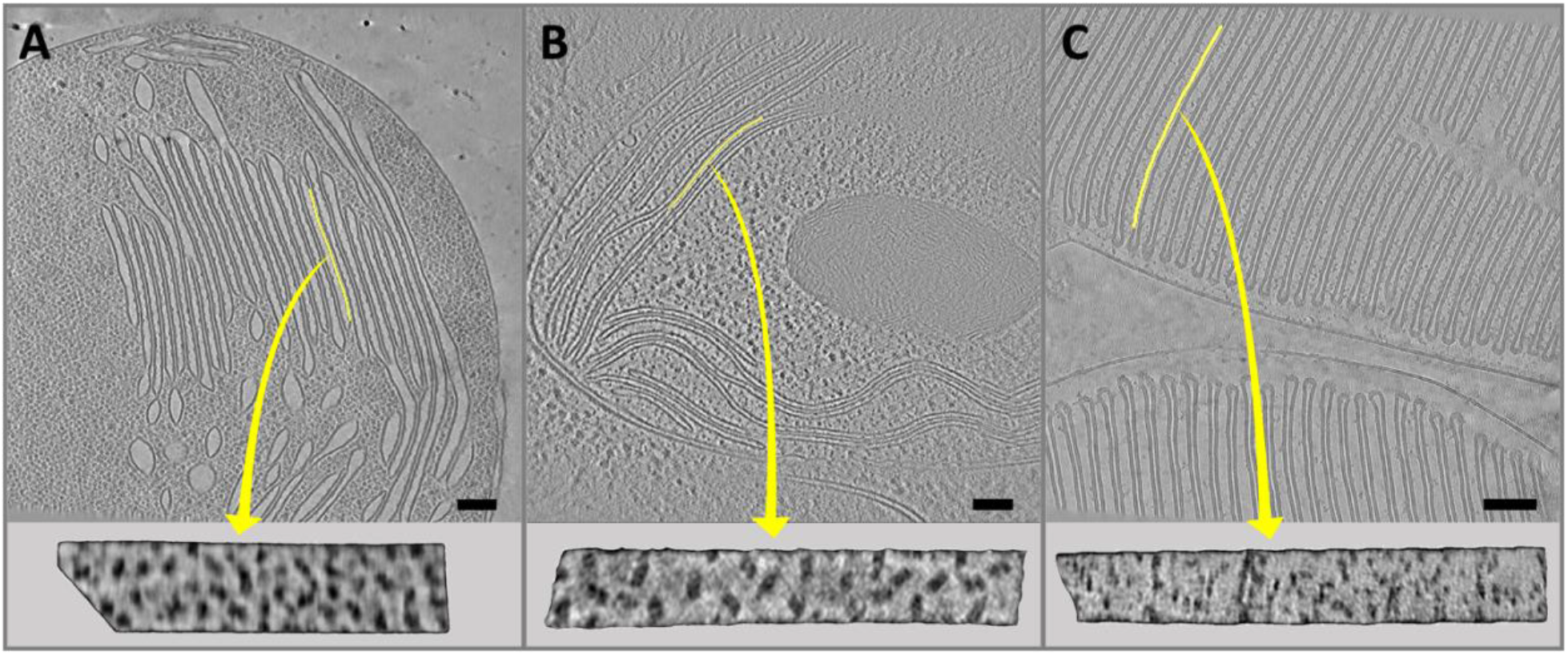
The three cryo-ET datasets examined in this study. For each panel, **Top:** Example 2D tomogram slices (scale bars 100nm), **Bottom:** Membranogram view of one membrane. The visualized membrane is highlighted in yellow. **A:** Dataset 1 – Chloroplasts isolated from *Spinacia oleracea*, acquired using defocus imaging and denoised with Cryo-CARE [25]. **B:** Dataset 2 [12] – Chloroplasts inside *Chlamydomonas reinhardtii* cells, acquired using Volta phase plate imaging [31], not denoised. **C:** Dataset 3 [26] – Rod outer segments isolated from wild-type mice, acquired using defocus imaging and denoised with Cryo-CARE.

### 3.2. Analysis of MemBrain’s Performance

#### Evaluation Metric

For evaluating MemBrain’s performance with respect to ground truth particle positions, we compute precision (*P*), recall (*R*), and F1 score (*F*_1_) of our predicted particle coordinates. For the calculation of recall, we define a true positive (*TP*_*R*_) as a ground truth (GT) PSII position that has been *hit*, i.e., a predicted position is within a certain radius (in this case, 4.5 voxels). Correspondingly, a false negative (*FN*_*R*_) is a GT position that is not hit by a predicted position. For precision, a true positive (*TP*_*P*_) is a predicted position that hits a GT position (either PSII or unknown densities), and a false positive (*FP*_*P*_) is a predicted position that does not hit a GT position. The F1 score is computed as the harmonic mean of recall and precision:

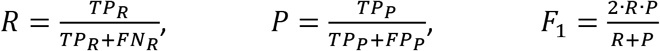

#### Quantitative Analysis

Table 1 shows the performance of MemBrain in comparison with other state-of-the-art deep learning protein picking algorithms, as well as the commonly used template matching approach (for more details about the generation of comparison positions, see supplementary S1.3). In addition, Figure 3 shows how well the methods performed with respect to ground truth on a 2D slice (top views), as well as two membranes (bottom views). All deep learning-methods were trained using 28 membranes from Dataset 1 as the training set (7 membranes were used for the validation set), while template matching was performed using a native structure of PSII embedded in a membrane density as the reference template. For evaluation, we only considered those predictions that were close to an annotated membrane in order to enable comparison to the corresponding ground truth. MemBrain outperformed the other methods on the test set of Dataset 1 and even generalized well to Dataset 2, despite a considerable domain shift between these samples. In particular, we emphasize that the recall for MemBrain far exceeded the level of the other methods, which is a critical advance to prevent missing ground-truth particles. In comparison, precision is less important, as there are mature techniques to clean up false-positive picks, e.g., via subtomogram classification [27]. Although DeepFinder picks well in some regions, it misses proteins in other regions completely (Figure 3, membrane views), conceivably because the training data does not cover all possible membrane orientations. EMAN already has a mechanism to compensate for the lack of particle orientations by performing random rotations on the training data. Nonetheless, it did not perform as well as MemBrain, demonstrating the importance of the subvolume rotation module and the focused detection in our pipeline. We conclude that all other methods have limitations in this very specific context of membrane proteins, as all of them require regions in the tomograms that are richly annotated, and expectedly have problems in this setting of very sparse labels. Compared to deep learning-based methods, template matching is even more difficult to tune, because the peaks in the detection are mostly triggered by the strong membrane signal and often do not correspond to actual protein complexes. As a result, template matching leads to low recall and precision. Besides particle detection, MemBrain can additionally extract particle orientations, which is beyond the capacities of other deep learning-based methods.

**Table 1.**
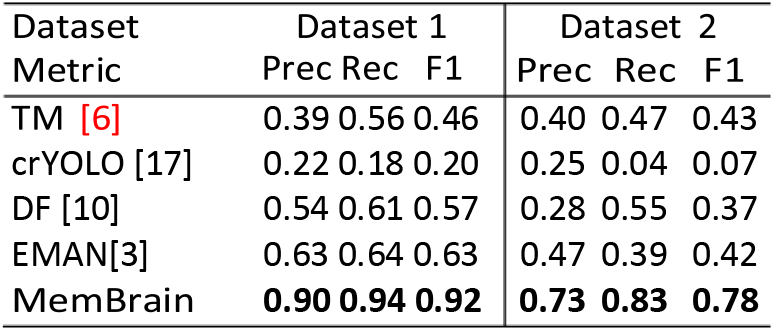
Benchmarking results on the test sets of Datasets 1 and 2 for Template Matching (TM) [29], crYOLO [9], DeepFinder [8], EMAN [7], and MemBrain.

| Dataset<br>Metric | Dataset 1 |  |  | Dataset 2 |  |  |
| --- | --- | --- | --- | --- | --- | --- |
|  | Prec | Rec | F1 | Prec | Rec | F1 |
| TM [6] | 0.39 | 0.56 | 0.46 | 0.40 | 0.47 | 0.43 |
| crYOLO [17] | 0.22 | 0.18 | 0.20 | 0.25 | 0.04 | 0.07 |
| DF [10] | 0.54 | 0.61 | 0.57 | 0.28 | 0.55 | 0.37 |
| EMAN[3] | 0.63 | 0.64 | 0.63 | 0.47 | 0.39 | 0.42 |
| MemBrain | **0.90** | **0.94** | **0.92** | **0.73** | **0.83** | **0.78** |

**Fig. 3.**
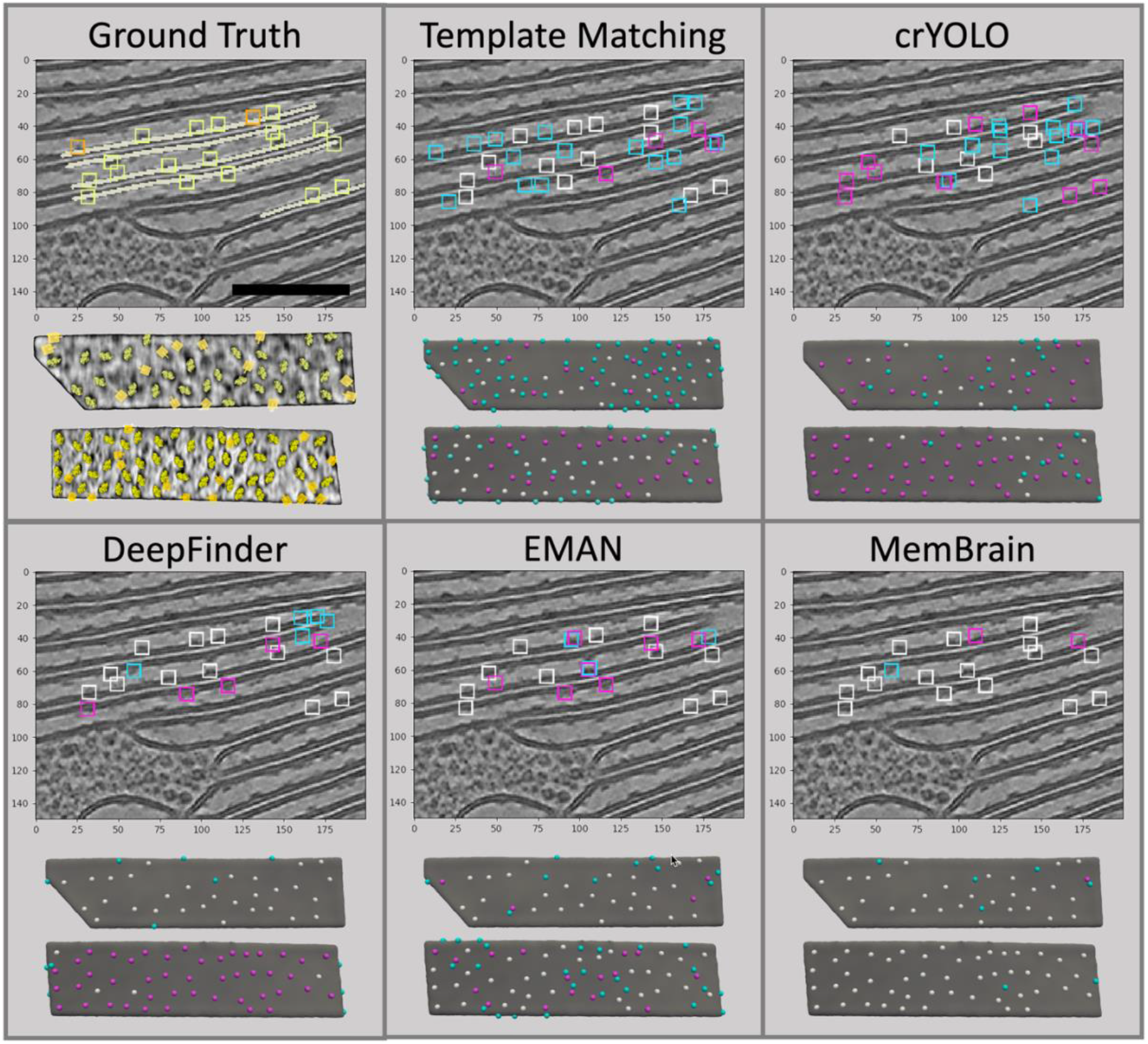
Visual analysis of predicted positions. For the ground truth panel, **Top:** 2D slice of a test set tomogram with annotated membranes highlighted in white (scale bar 100nm). Yellow boxes show PSII positions, orange boxes represent unknown density (UK) positions. **Bottom:** Membranogram views of two membranes from the test set. Mapped in are PSII protein structures (yellow) and UK positions (orange cubes). The five other panels show the analysis of predicted positions for all compared methods. True positives are highlighted in white, false negatives in magenta, and false positives in cyan.

Figure 4 shows the distribution of MemBrain’s predicted orientations with respect to the ground truth. We achieved a mean absolute error of 24.4 degrees (maximum possible deviation is 90 degrees due to PSII’s C2 symmetry) on our Dataset 1 test set, which can serve as a good initialization for subvolume alignment and make the subsequent subtomogram averaging [5,28,29] more efficient.

**Fig. 4.**
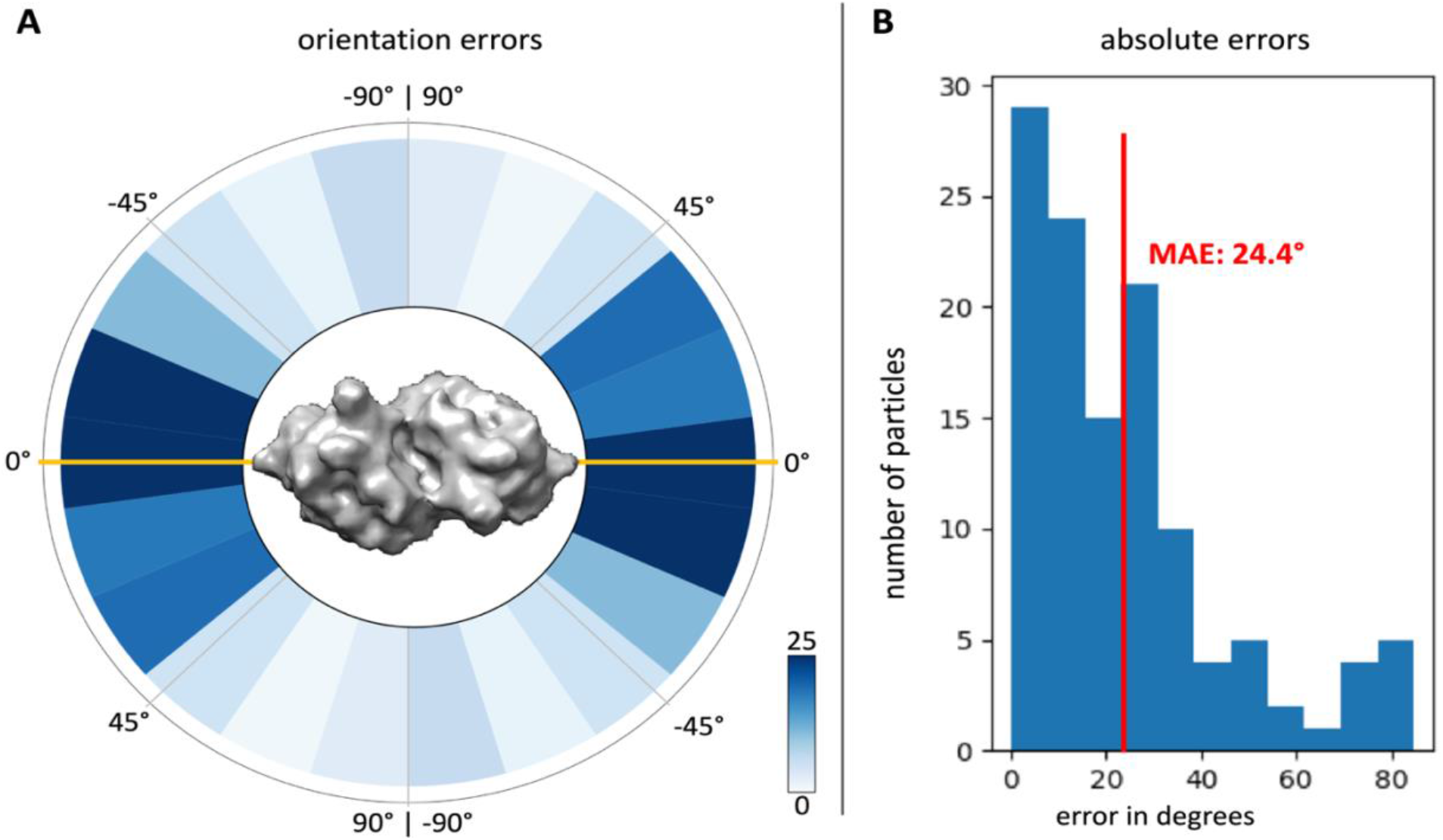
Analysis of particle orientations estimated by MemBrain. Comparison of MemBrain’s 120 true positive picks to the ground truth annotations in the Dataset 1 test set. **A:** Circular heatmap showing differences between ground truth in-plane angles and estimated angles. The long axis of the mapped PSII structure (0°) is plotted in orange. Note that due to PSII’s two-fold symmetry, the deviation cannot exceed 90°. **B:** Histogram of absolute differences between ground truth in-plane angles and estimated angles. The mean absolute error (MAE) is indicated in red.

#### Qualitative Evaluation on Dataset 3

For Dataset 3, we did not have ground truth protein positions to compute quantitative evaluation metrics. Instead, we assessed the picks from MemBrain and PySeg using subtomogram averaging (Figure 5; For details about the averaging procedure, see supplementary S.1.4). First, we trained a MemBrain model using the ground truth data from Dataset 1 (the same model as for the previous section), which reliably picked PSII complexes in this spinach dataset (Table 1; Figure 5A). Then, we applied this pre-trained MemBrain model to predict membrane-bound protein positions in the mouse rod outer segments of Dataset 3 (Figure 5B). For comparison, we also predicted membrane-bound protein positions in Dataset 3 using PySeg (Figure 5C). Subtomogram averaging of the MemBrain picks, PySeg picks, as well as intersection and difference sets between these picks revealed interesting differences between the two detection algorithms. In both Datasets 1 and 3, MemBrain consistently picks densities that are embedded in the membranes and appear as small bumps. In contrast, PySeg was configured to pick all densities over the membranes regardless their shape. This initial picking is refined after the first cleaning step, where the expert selected automatically generated 2D rotational average classes that indicated a density on the membrane (PySeg_clean_, see supplementary S.1.2, and Figure S7). Thus, PySeg picks a wider variety of particles, resulting in a different average. Looking at the consensus analysis (Figure 5D), we can see that PySeg picks often coincide with MemBrain picks, and the average of this intersection subclass looks very similar to the pure MemBrain average. In the subclass of MemBrain picks that were missed by PySeg, we see a smaller bump on the membrane. On the other hand, the PySeg picks that MemBrain missed appear to be densities that are suspended above the membrane and not embedded within it. A further classification of this subclass can be seen in Figure S9. It is worth noting that this behavior of MemBrain is often desirable, since it has been trained to find exactly these bumps on the membrane (see the PSII average in the top row of Figure 5), and one may not want to detect everything that is close to a membrane, but rather only specific membrane-embedded proteins. PySeg is a comprehensive data driven workflow, but a specialized pipeline like MemBrain can save a lot of time and effort when the membrane proteins are visually recognizable.

**Fig. 5.**
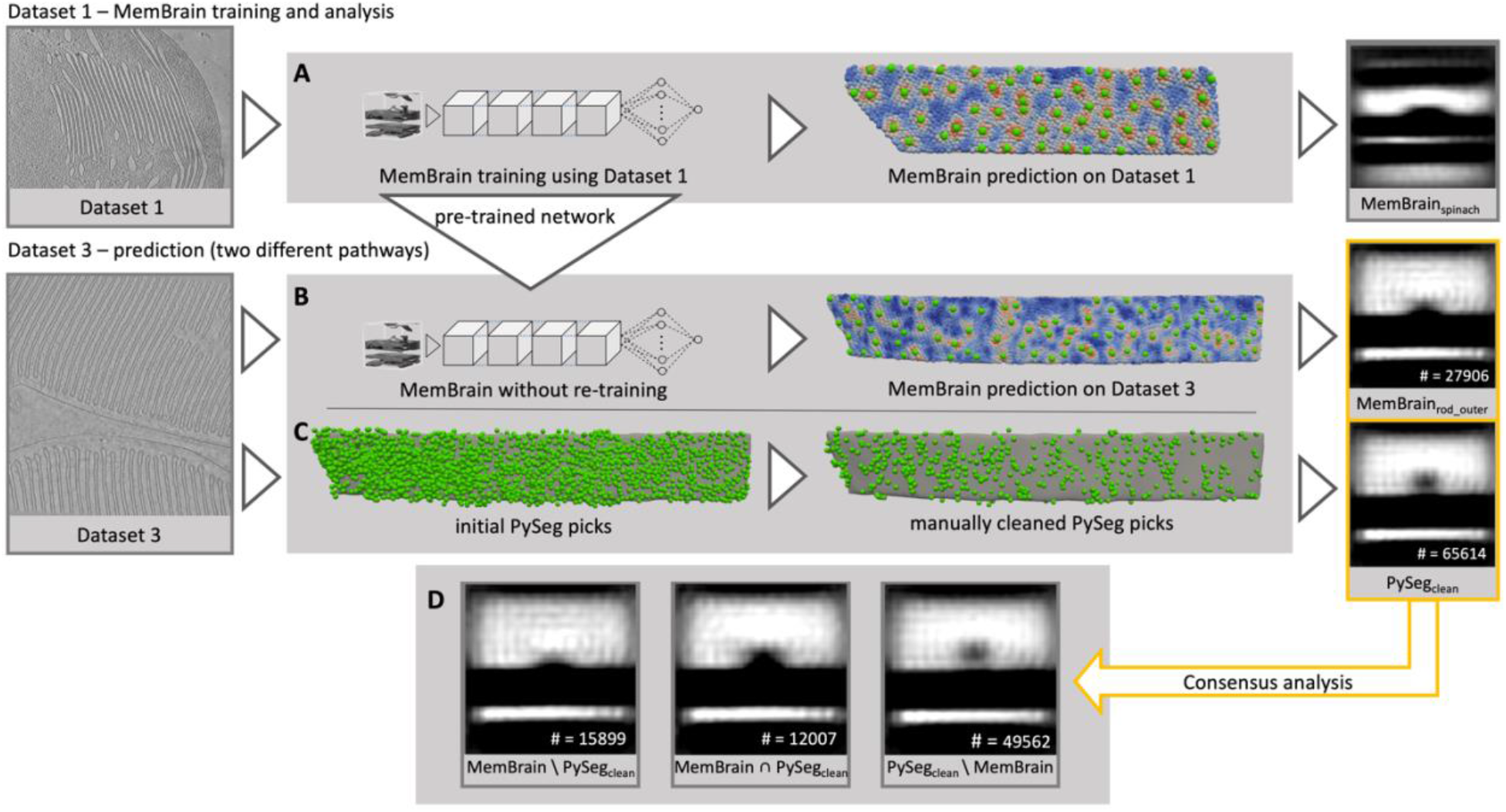
Qualitative comparison of MemBrain and PySeg on Dataset 3. **A:** MemBrain is trained using the ground truth data from Dataset 1. The trained program is used to predict particle positions (green) in the test set of Dataset 1. **B:** The same MemBrain program (trained on Dataset 1) is used to predict particle positions (green) in Dataset 3. **C:** PySeg is applied to generate particle positions (green) in Dataset 3. Visualized are the initial picks of PySeg (left) and the picks after cleaning using selection of 2D rotational averages of PySeg (right). Rotationally symmetrized subtomogram averages were generated from the picks of each pathway (right column). **D:** Consensus analysis using subtomogram averages to compare the MemBrain picks and the cleaned PySeg picks. MemBrain \ PySeg_clean_ = positions picked by MemBrain but not PySeg_clean_. MemBrain ∩ PySeg_clean_ = positions picked by both MemBrain and PySeg_clean_. PySeg_clean_ \ MemBrain = positions picked by PySeg_clean_ but not MemBrain.

#### Ablation Study

In our ablation study (Table 2), we show the results of MemBrain with varying experimental settings. First, we explored the minimum amount of training data that is required to train our model. Like for most biomedical images, data annotation requires human expertise and is generally difficult to obtain. In our experiment, we used 1, 8 or 28 annotated membranes in the training set, and we performed hyperparameter tuning using grid search for both learning rate (range 3×10^−5^ to 3×10^−4^) and weight decay parameters (range 0 to 1×10^−3^) although our pipeline was quite robust with respect to these parameters. The models were trained using a batch size of 1024 for up to 5000 epochs, with early termination in case the validation stopped decreasing. As shown in Table 2, even with only one annotated membrane, MemBrain still achieved an F1 score of 0.88, which far exceeded EMAN, DeepFinder, and crYOLO (Table 1), even though these other deep learning networks were trained on the entire training dataset of 28 membranes. Furthermore, we explore the contribution of individual components in the MemBrain pipeline. We find that denoising the tomograms is vital to ensure a good performance, as it simplifies the task by removing the necessity for the model to learn to cope with noise. In addition, using a CNN for a regression task rather than more commonly-used classification also improves the overall performance of the entire pipeline, particularly when the training dataset is small. We reason that the induced label noise from inaccurate manual labeling of particle center positions has a larger influence on classification than on distance regression, where the score assignment is smoother. Finally, we explored the influence of our subvolume normalization module: The values for “no normalization” correspond to the results for MemBrain without normalized volumes, but trained with arbitrary subvolume rotations as data augmentation during training. Again, especially for the one membrane (1 mb) and eight membranes (8 mbs) settings, the results are worse than our fully trained MemBrain model. This corroborates the importance of our normalization module for simplifying the detection problem.

**Table 2.** Ablation study: Effects of using a regression target (as compared to classification), denoised data, and the rotational normalization module for 1, 8, and 28 membranes, respectively.

| Method | 1 mb | F1 Score |  |
| --- | --- | --- | --- |
|  |  | 8 mbs | 28 mbs |
| Full MemBrain | 0.88 | **0.91** | **0.92** |
| Classification | 0.79 | 0.80 | **0.91** |
| Non-denoised | 0.78 | 0.80 | 0.85 |
| No normalization | 0.81 | 0.86 | 0.87 |

## 4 Conclusion

In this study, we present MemBrain, a deep learning-based pipeline that is specialized for automatic detection of membrane-bound proteins in cryo-electron tomograms. In the preprocessing step, MemBrain uses the membrane geometry and rotates tomographic subvolumes into a normalized orientation, which generalizes the trained CNN to membranes with different orientations than the training set. Moreover, this rotation step reduces the complexity of the protein detection task, and thus, our network can be efficiently trained with a small dataset consisting of only one annotated membrane. This is experimentally practical, as cryo-ET data is time-consuming to generate, and the manual annotation is laborious. We evaluated MemBrain on three different datasets and showed that our pipeline outperforms other state-of-the-art methods (both conventional and deep learning-based) for particle picking by a large margin (recall: MemBrain >0.9 vs. other methods <0.65). In Dataset 3, where no ground truth annotations were available, we showed qualitatively that MemBrain’s picks resemble membrane-embedded proteins. Furthermore, MemBrain is able to estimate protein orientations, which is beyond the current capacities of other deep learning methods. The particle positions and orientations extracted by MemBrain can be used to analyze the organization of protein complexes within membranes, revealing how they interact with each other to drive cellular processes. Since MemBrain is generalizable to unseen data domains, it will likely be applicable to investigating biological mechanisms in various kinds of membranes, ranging from protein production in the endoplasmic reticulum to the bioenergetic reactions in mitochondrial cristae and chloroplast thylakoids.

## 5 Acknowledgements

Calculations were performed at the Max Planck Institute for Biochemistry computing cluster. We thank Matt Johnson for collaboration in producing Dataset 1. L.L. acknowledges funding from the Munich School for Data Science (MUDS) and a fellowship from the Boehringer Ingelheim Fonds. R.D.R. acknowledges funding from the Alexander von Humboldt Foundation and a non-stipendiary fellowship from EMBO. W.W., T.P. and B.D.E. were supported by DFG grant no. EN 1194/1-1 as part of FOR 2092 and the Helmholtz Association. A.M.-S. was supported by the Deutsche Forschungsgemeinschaft (DFG, German Research Foundation) under Germany’s Excellence Strategy EXC 2067/1-390729940. The authors declare no conflict of interest.

## Supplementary information

### 1.1 Description of datasets

#### Datasets 1 and 2

Cells and chloroplasts were plunge frozen using a Vitrobot Mark 4 (FEI Thermo Fisher) and stored in liquid nitrogen. Cryo-FIB milling was performed using a Quanta dual-beam FIB/SEM (*Chlamydomonas*, Dataset 2) or Aquilos dual-beam FIB/SEM (Spinach, Dataset 1) instrument (FEI, Thermo Fisher Scientific), as previously described[24]. Tomographic data was acquired on a 300 kV Titan Krios microscope (FEI, Thermo Fisher Scientific), equipped with a post-column energy filter (Quantum, Gatan) and a direct detector camera (K2 Summit, Gatan). Dose-symmetric (Dataset 1) or bidirectional tilt-series (Dataset 2) were acquired with SerialEM software [30] using 2° steps between −60° and +60°. Individual tilts were recorded in movie mode with 12 frames per second, at an image pixel size of 3.42Å (42000x magnification). The total accumulated dose for the tilt-series was kept around 100 e^−^/Å^2^. The *Chlamydomonas* tilt-series (Dataset 2) [12] were acquired with a Volta Phase Plate at −0.5 μm defocus [31], while the spinach tilt-series (Dataset 1) were acquired with a standard objective aperture and defocus values from −4 to −4,5 μm. Using IMOD software [32], tilt-series were aligned with patch tracking and reconstructed with weighted back projection. Additionally, Dataset 1 was dose weighted with TOMOMAN software [33] and denoised with cryo-CARE [25].

#### Dataset 3

Rod outer segments were isolated, vitrified on EM grids, thinned by cryo-FIB milling, and imaged by cryo-ET as described in [26]. Tilt-series were acquired on the same Titan Krios microscopeas Datasets 1 and 2. Bidirectional tilt-series were collected between +50° and −50°, starting at 20° with a tilt increment of 2° and a total dose of 100 e^−^/Å^2^. The individual projection images were recorded at an image pixel size of 2.62 Å (53000x magnification) in movie mode with 6 frames per tilt at tilt angle α = 0°. The number of frames per tilt were increased at higher tilts according to the 1/cos(α) as implemented in SerialEM. Standard objective aperture imaging was used, with a target defocus of −3 μm. The acquired tilt-series were aligned using Platinum particles on the lamella surface as fiducial markers [26] and reconstructed in IMOD software. Furthermore, dose weighted was performed with TOMOMAN software and denoising with cryo-CARE similar to Dataset 1.

**Fig. 6. SUPPLEMENTARY FIGURE.**
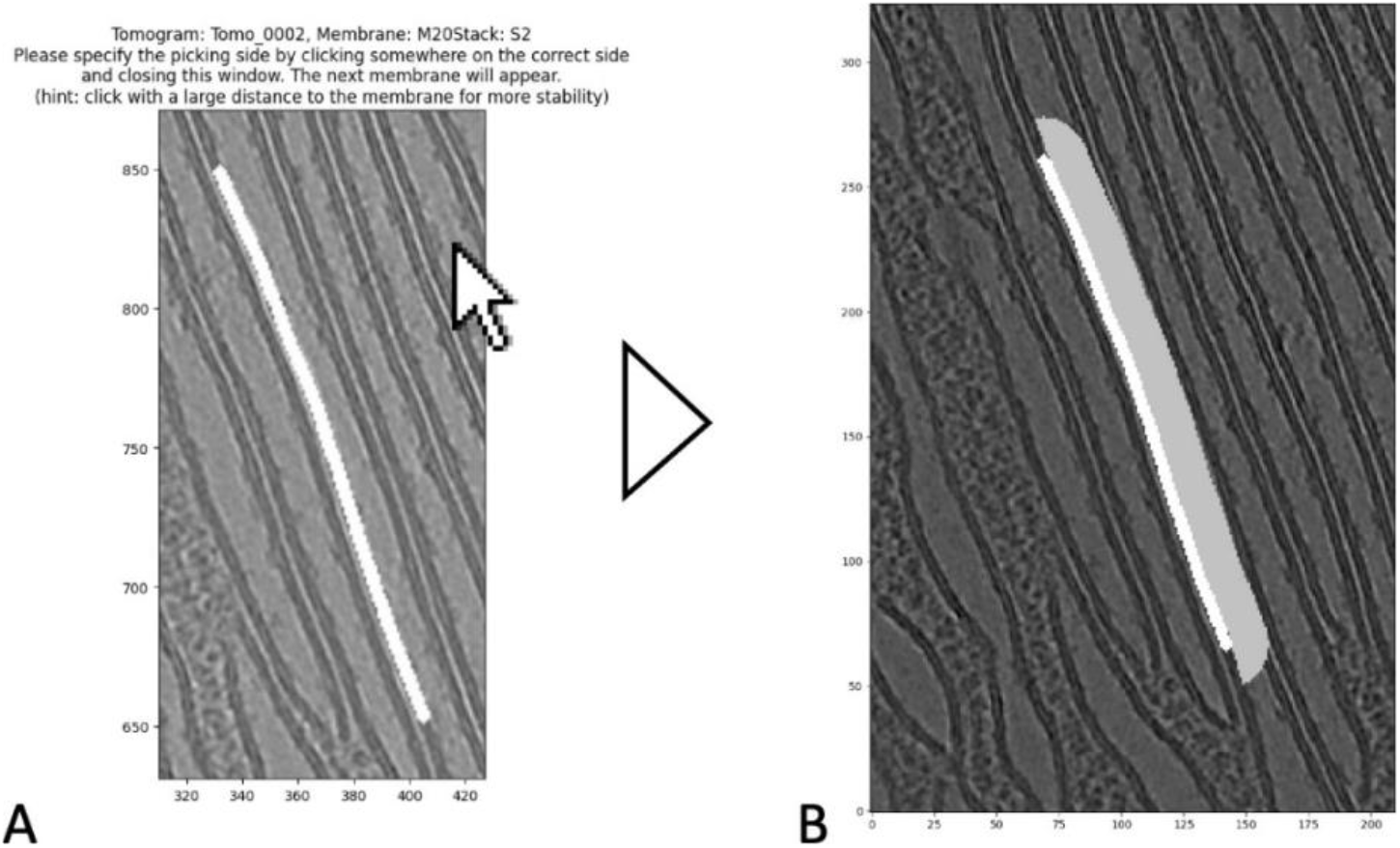
Graphical assistance for using MemBrain. White segmentations represent given membrane segmentations. A: Membrane selection interface. The user should click on the side of the membrane where they want to detect proteins. B: Membrane segmentation and resulting selected side of the membrane (grey: thresholded neighborhood used to estimate normal vectors), visualized using MemBrain’s membrane side inspection script, which can be used to verify that the correct sides were picked.

**Fig. S7. SUPPLEMENTARY FIGURE.**
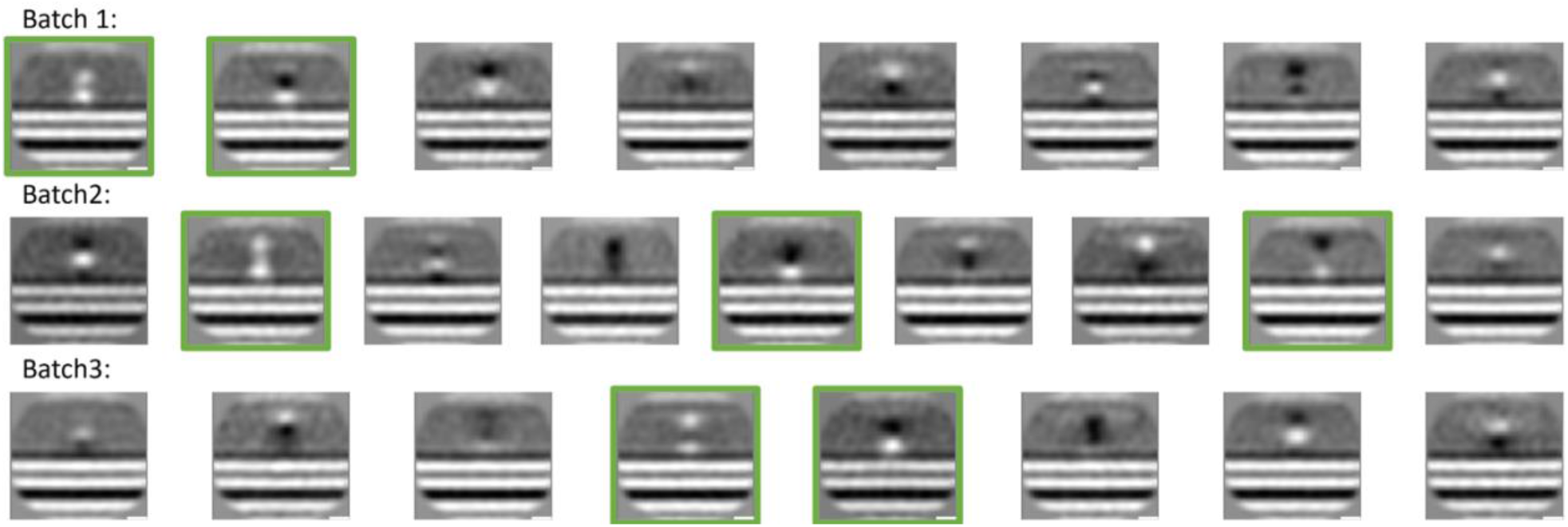
Selection of 2D rotational average classes. 270k particles were extracted, split into batches of 30k particles each, and clustered into several classes using PySeg’s 2D rotational average clustering. From the resulting classes, those with a density in contact with the membrane were selected (green).

### 1.2 PySeg protein picking

Protein picking in PySeg was performed using the default parameters. Only the vertex density during the calculation of the graphs was set to 0.01 per nm, a value much higher than expected, to assure that all possible densities are picked. The minimum allowed distance between adjacent proteins was set to 4 nm refinement of the membrane normal vectors and the alignment of the membranes was done in STOPGAP (v.0.7.1) [29]. The star-file as output of PySeg protein picking was converted into a STOPGAP motivelist, subvolumes were extracted from 4x binned non-CTF-corrected tomograms (box size = 64 voxel, voxel size = 1.048 nm) and aligned with shifts only allowed perpendicular to the membrane. Subvolume were re-extracted at the refined positions with a smaller box size of 32 voxels and the refined motivelist converted into a star-file for PySeg. Afterwards, PySeg’s plane_align_class.py script was used for two successive rounds of 2D classification (Affinity propagation, preference = −10, pp 3D = False). Here, a mollified cylinder mask (radius = 10 voxel, height = 10 voxel) focused the classification on the cytosol adjacent to the membrane without including the membrane signal itself. In the first round (referred to as PySeg_clean_ in the manuscript), all picked proteins were processed in batches of 30k subvolumes and those classes that indicated a density in contact with the membrane irrespective of its size were selected (see Figure S7). In a second round, all selected proteins were classified together and only those classes with reasonably large density were selected (these picks were not used for the evaluation in this paper).

### 1.3 Configurations of comparison methods

#### DeepFinder

Ground truth protein positions and orientations were used to map a thresholded volumetric model of PSII into the tomograms. This gave us an accurate segmentation mask, where, for all training protein positions, all voxels that are occupied by a protein density are labeled as 1. The remaining voxels are labeled as background (i.e., 0). We trained DeepFinder using the same data split as we did for MemBrain (28 membranes training set, 7 validation set). In addition to DeepFinder’s default settings, we also performed hyper-parameter tuning for learning rate and random shifts range, and found the best performance on the validation set for a learning rate of 1 × 10^−4^, and random shifts in the range of 11.

As preprocessing for our tomograms, we used the default preprocessing suggested in EMAN: low- and high-pass filtering, followed by normalization (to mean 0 with standard deviation 1) of the tomogram. Finally, values are clamped between −3 and 3 to remove outliers.

#### EMAN (v.2.91)

We used the same ground truth segmentation masks from DeepFinder to generate the ground truth for EMAN. Because EMAN required annotated 2D slices, we extracted 2D slices (size 64 x 64) from the 3D annotated volumes, each centered around a protein position to ensure that the patches contain proteins. We performed the default preprocessing (as described in the DeepFinder section), and also did hyperparameter tuning. EMAN’s architecture is very shallow, which can avoid overfitting on the training data. Nevertheless, we wanted to rely on more information than only the training loss. So, we trained several models to the end, and then performed tuning of the extraction parameters on the validation set to find the best models and setting: we trained models for varying numbers of epochs, and different learning rates. Then, for each model, we tuned the protein extraction parameters for the density threshold, and the mass threshold (size of an object to be considered a true object). Here, we found the best model with a learning rate of 1 × 10^−5^, trained for 5000 epochs, and, for the extraction, used a mass threshold of 2.0, and a density threshold of 0.5.

#### crYOLO (v.1.8.0)

Instead of volumetric segmentation masks, crYOLO requires only the positions of the ground truth particles. Therefore, we converted our manually picked positions into the required format. crYOLO requires the user to label a protein in a 2D slice, even if only a small portion of it can be seen. Therefore, we experimented with transferring ground truth positions also to adjacent slices, but did not find an improvement in performance. Again, we performed hyper-parameter tuning. For training the neural network, we tuned the parameter “object_scale”, determining the weight of true positive samples in the loss function (compared to the weight of false positives). Furthermore, we tuned parameters for the extraction of protein positions: The threshold for the network-assigned scores, the minimum distance between boxes, and the minimum number of boxes in one trace to be considered a protein. For us, an object scale of 50 (very high, probably necessary due to the sparse annotations), a score threshold of 0.05, a minimum distance of 5, and a minimum number of boxes in a trace of 2, were the best settings.

#### Template matching

Template matching experiments were performed in STOPGAP [29]. A previously obtained average of PSII [12] was used as template for detection of this protein in Datasets 1 and 2 using an angular search step of 20 degrees in bin4 tomograms (pixel size: 13.68 Å). Wedge lists containing information on electron exposure, defocus and tilt angle of the tilt series were obtained from the IMOD reconstruction metadata [32] for each tomogram. This information is used to properly weight the Fourier transform of the template rotated in each orientation and constrain the cross-correlation only to the areas where information is present. To avoid overfitting, the template was low-pass filtered to ~40 Å during search. For performance analysis, we visually tuned the number of extracted picks, as well as the minimum distance between two picks. We arrived at values of 400 picks per tomogram, and a minimum distance of 13 voxels, giving the best performance on the test tomograms.

### 1.4 Subtomogram averaging for Dataset 3

To create the averages in Figure 5 and Figure S9, we utilized the following procedure: We extracted protein positions using MemBrain and compared them to the ones obtained by PySeg (see section S.1.2). We subdivided the picks into six subsets, as can be seen in Table S3. For subsets 1, 3, 4, and 5, we used the normal voting procedure described in Section 2.2. to receive membrane normal vectors for each position. Then, from the normal vectors, we computed the Euler angles that describe the rotation to align the membrane horizontally. For subsets 2 and 6, we used the Euler angles that are computed by PySeg. Then, we used STOPGAP [29] to create averages of the picked positions in bin4 (pixel size: 10.48Å). First, we created an initial average. Afterwards, we refined this average in 3 rounds of subtomogram alignment, with the goal of refining the membrane alignment. For this, we used a cylindrical mask to focus the alignment only on the current membrane and the density on top of it. We allowed only small shifts in x-, y, and z-direction (2 bin4 voxels in each direction), and used a strong symmetry constraint (C20). For subset 6, we further performed subtomogram classification into 3 classes using STOPGAP’s “ali_multiref” function. We performed this classification step for 10 rounds in order to investigate if there is a class corresponding mainly to membrane-embedded proteins.

**Table S3.** Overview of different subsets of Dataset 3 picks for averaging comparison. “Round 1” corresponds to the first PySeg pick cleaning step, where only averages containing membranes with densities were selected.

| Subset name | Description | # positions |
| --- | --- | --- |
| MemBrain<sub>rod_outer</sub> | All MemBrain picks | 27906 |
| $\text{PySeg}_{\text{clean}}$ | All PySeg picks after round 1 | 65614 |
| $\text{MemBrain} \setminus \text{PySeg}_{\text{all}}$ | MemBrain picks, missed by PySeg before round 1 | 2163 |
| $\text{MemBrain} \setminus \text{PySeg}_{\text{clean}}$ | MemBrain picks, missed by PySeg after round 1 | 15899 |
| $\text{MemBrain} \cap \text{PySeg}_{\text{clean}}$ | MemBrain picks, coinciding with PySeg picks after round 1 | 12007 |
| $\text{PySeg}_{\text{clean}} \setminus \text{MemBrain}$ | PySeg picks after round 1, missed by MemBrain | 49562 |
| $\text{PySeg}_{\text{clean}} \setminus \text{MemBrain}$<br>Class 1 | Classified using subtomogram classification in STOPGAP | 9665 |
| $\text{PySeg}_{\text{clean}} \setminus \text{MemBrain}$<br>Class 2 | Classified using subtomogram classification in STOPGAP | 21678 |
| $\text{PySeg}_{\text{clean}} \setminus \text{MemBrain}$<br>Class 3 | Classified using subtomogram classification in STOPGAP | 18219 |

**Table S4.** Overview of our network architecture. Conv=Convolutional layer; BN=Batch normalization

| Layer | Type | # Filters | Size |
| --- | --- | --- | --- |
| 1 | Conv (valid) | 32 | 3x3x3 |
|  | BN |  |  |
|  | ReLU |  |  |
| 2 | Conv (same) | 32 | 3x3x3 |
|  | BN |  |  |
|  | ReLU |  |  |
| 3 | Conv (valid) | 32 | 3x3x3 |
|  | BN |  |  |
|  | ReLU |  |  |
| 4 | Conv (same) | 32 | 3x3x3 |
|  | BN |  |  |
|  | ReLU |  |  |
| 5 | Linear | 1 | $(\text{box\_size} - 4)^3 * 32$ |

**Fig. S8. SUPPLEMENTARY FIGURE.**
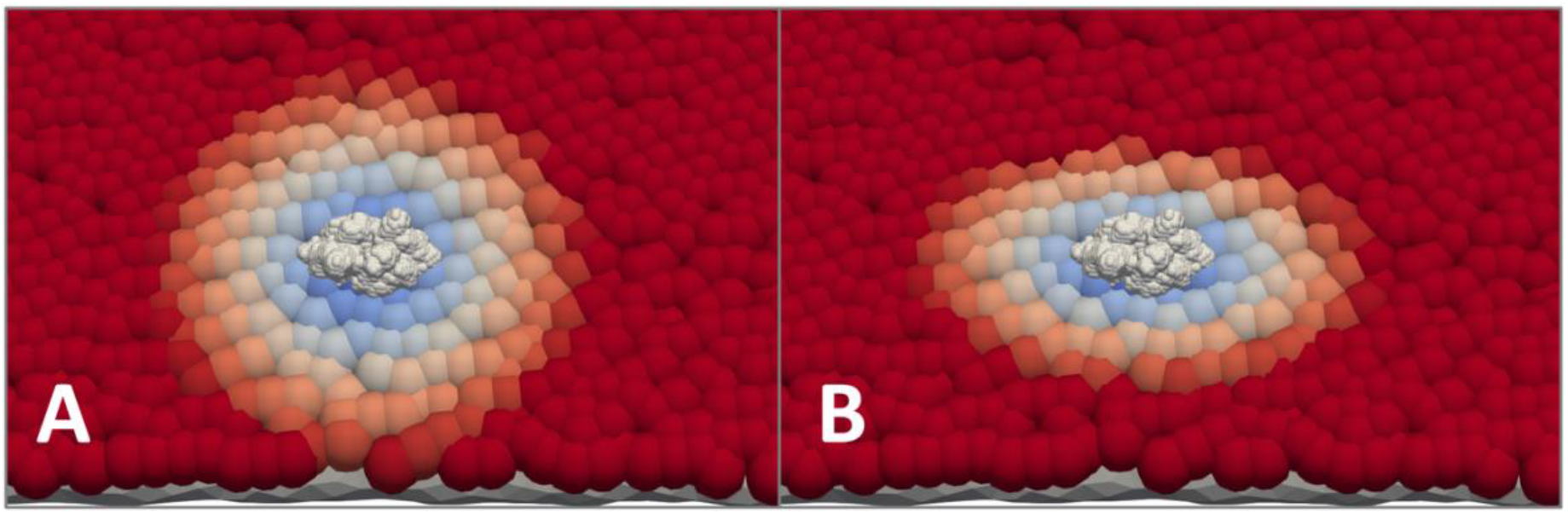
Visualization of the score assignment using the Mahalanobis distance. The white density represents a PSII particle, mapped into its ground truth position and orientation. Distance values are represented by colors: blue points correspond to points with low distances. Red points have a higher assigned distance. A: Visualization of the Euclidean distance. B: Visualization of the Mahalanobis distance.

**Fig. S9. SUPPLEMENTARY FIGURE.**
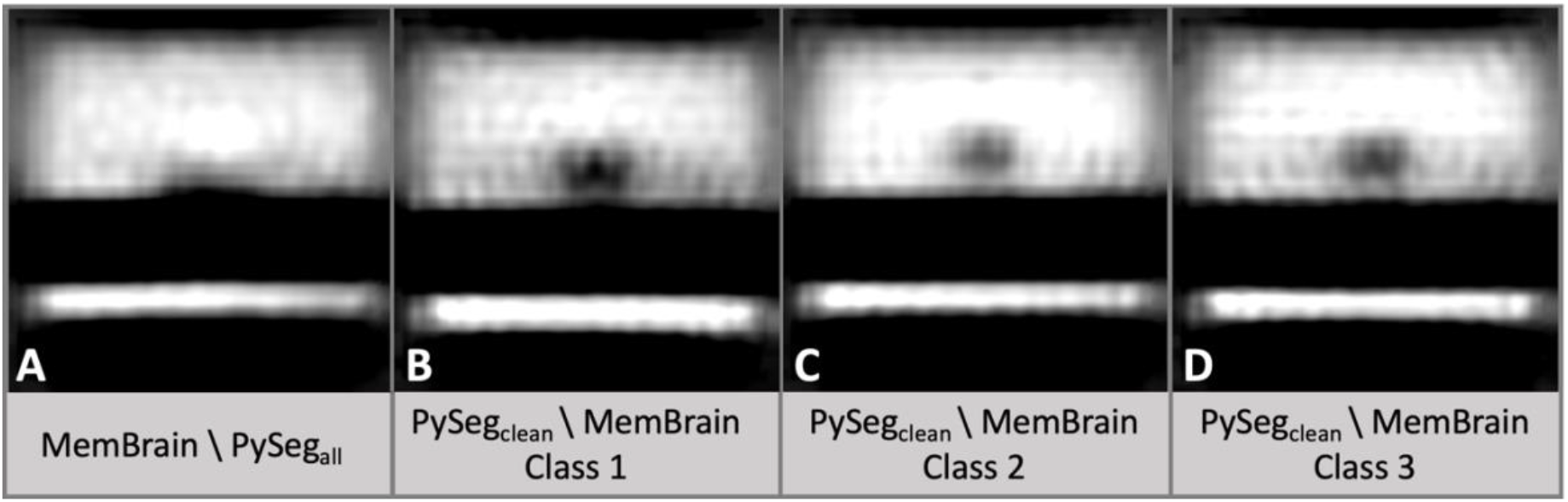
Rotationally symmetrized subtomogram averages generated from different subset of picked positions from Dataset 3. A: MemBrain positions that were missed by PySeg (before cleaning steps). B-C: Subtomogram classification results of (PySeg_clean_ \ MemBrain) into 3 classes. For more details on how the averages were created, see S.1.4. and Table S3.

